# Instruction-tuned extraction of virus-host interactions from integrated scientific evidence

**DOI:** 10.1101/2025.09.02.673691

**Authors:** Zheng Zhang, Weichao Li, Jiajun Ren, Yufei Chen, Ben He, Qiang Sun, Haibo Wang

## Abstract

**Motivation:** Viral infectious diseases continue to pose a major threat to global health. Understanding protein-protein interactions (PPIs) and RNA-protein interactions (RPIs) between viruses and hosts is essential for elucidating infection mechanisms. However, manual curation of these interactions from biomedical literature is inefficient, creating a pressing need for automated and scalable extraction methods. Large language models (LLMs), such as the generative pre-trained transformer (GPT) and bidirectional encoder representations from transformers (BERT), offer promising solutions. Yet, most existing datasets focus on abstracts, overlooking other information-rich sections. We aim to develop a data-efficient approach to extract virus-host interaction (VHI) entities from full-text biomedical articles, including Results, Methods and tables. To our knowledge, this is the first study to apply instruction tuning to full-text VHI extraction

**Results:** We curated a dataset containing 3, 395 PPI and 674 RPI entities from the Results, Materials and Methods sections, along with 566 PPIs and 793 RPIs from tables. Under low-resource conditions (<500 training examples), our instruction-tuned ChatMed-VHI model achieves the best overall performance (F1: 89.7%, Precision: 95.3%), outperforming PubMedBERT (F1: 74.6%, Precision: 75.1%). When scaled to the full dataset (>4, 000 training examples), ChatMed-VHI maintained the highest overall performance, while PubMedBERT achieved slightly higher precision (92.3% vs. 91.3%). Notably, ChatMed-VHI improved F1 and recall by 2.79% but precision dropped by 4.20% with more training data, whereas PubMedBERT improved consistently across all metrics. These results demonstrate the effectiveness of instruction-tuned LLMs for full-text biomedical extraction tasks, and position ChatMed-VHI as a scalable, domain-adaptable solution for VHI mining.

## Introduction

Virus-host interactomes encompass the intricate molecular mechanisms underlying viral infections, including Protein-Protein Interactions (PPIs), RNA-Protein Interactions (RPIs), RNA-RNA Interactions (RRIs), and DNA-Protein Interactions (DPIs). Understanding these interactions is critical for deciphering infection pathways and immune responses. To support such investigations, a variety of interaction databases have been established. General PPI databases such as BioGRID [1], IntAct [2], and STRING v11 [3] cover a wide range of intra- and inter-species protein-protein interactions. In contrast, specialized resources such as HVIDB [4], HVPPI [5] and VirusMentha [6] focus specifically on virus-human protein interactions, while VIRbase [7] and CoVInter [8] curate viral RNA-host interactions, particularly those involving RNA-binding proteins. Despite the availability of curated databases, manually extracting virus-host interaction data from literature remains a formidable challenge. Manual data curation is not only time-consuming but also demands expert knowledge of virology, host biology, molecular entities (e.g., RNAs and proteins), and experimental methodologies [9]. To address this bottleneck, researchers have increasingly turned to NLP-based extraction methods. However, most existing benchmarks and applications rely solely on abstracts, ignoring the majority of structured and contextual information present in full-text documents.

This limitation has been well documented. For example, Islamaj et al. (2021) showed that chemical NER systems trained only on titles and abstracts suffer substantial performance drops when applied to full texts, which contain more complex language but also richer information, such as compound properties, biological effects, and inter-entity interactions. In their analysis, over 70% of unique MeSH assignments and 78% of distinct chemical mentions,including synonyms and variants,were found only in full text [10]. Similarly, Yeganova et al. (2021) found that different full-text sections contribute unequally to retrieval performance, and that section-aware scoring models outperformed abstract-based ones on biomedical datasets [11]. Beyond individual sections, relevant information is often dispersed across the entire article, including narrative (e.g., Results, Methods) and non-narrative (e.g., tables, supplementary materials) formats. In virus-host interaction related articles. Materials and Methods sections often contain key contextual information, such as virus strains, host cell types, tissue origins, and infection durations, which are typically omitted from abstracts. The Results section may describe important PPIs and RPIs involved in viral infections or immune evasion, while tables often present large amounts of interaction data not explicitly mentioned in the main text. This heterogeneity in data types and formats hinders scalability, underscoring the need for an automated and integrative extraction framework capable of handling the full scope of scientific articles.

Recently, prompt-based frameworks such as ChatExtract [12] has demonstrated the feasibility of applying conversational LLMs directly to full-text data extraction. ChatExtract uses a set of engineered prompts to guide models like GPT-4 in identifying and validating data points across entire documents, achieving high precision and recall in materials science tasks. Its performance highlights the potential of LLMs to process long-length scientific content [12, 13]. While effective for structured data, it lacks the ability to capture complex biomedical interactions, such as virus-host PPIs and RPIs, which are often expressed across multiple sentences and sections of full-text articles.

In parallel, advances in biomedical natural language processing (BioNLP) introduce powerful pretrained models fine-tuned for specific downstream tasks [14]. For example, Bidirectional Encoder Representations from Transformers (BERT) [15] pretrained on large general-domain corpora achieved remarkable performance on a series of NLP tasks including named entity recognition (NER) [16], relation extraction [17], document classification [18], and question answering [19]. BERT-based variants such as PubMedBERT [20], BioBERT [21], and BioLinkBERT [22] have further improved performance on biomedical NER tasks. In particular, BioLinkBERT incorporated document connections during pretraining [22], and outperformed PubMedBERT in the BLURB benchmark dataset [23]. However, these BERT-based struggle with recognizing non-contiguous or complex biomedical entities that are dispersed throughout the text [24]. Second, they tend to perform poorly in low-resource scenarios where limited training data is available, hampering their ability to generalize across domains [25]. Third, the integration of multiple small datasets often introduces inconsistencies in annotation standards, such as variations in entity types, span boundaries, and normalization, ultimately diminishing model performance [26].To overcome these limitations, recent research has shifted towards text-generative large language models (LLMs), such as the GPT series and LLaMA, offering greater flexibility than BERT-based models through prompt engineering and instruction tuning [27]. While early evaluations revealed that ChatGPT achieved competitive results on question answering tasks compared with BERT-based models, its performance was suboptimal in the absence of domain-specific fine-tuning [23]. Similarly, comparative studies suggested that GPT-based models underperformed relative to BioBERT in PPIs classifications[28]. In response to these challenges, BioInstruct, an instruction-tuned variant of LLaMA, demonstrated strong performance across various biomedical information extraction tasks [29]. Keloth et al. applied instruction tuning to LLaMA-7B using domain-specific prompts derived from biomedical NER datasets, consistently outperforming few-shot GPT-4 and matching or exceeding PubMedBERT on multiple benchmarks [30]. These instruction-tuned LLMs demonstrated promising results in low-resource and cross-domain BioNLP applications.

While these LLMs developments are encouraging, biomedical information extraction tasks such as virus-host interaction (VHI) extraction require not only flexible LLMs but also section-aware and cross-sentence understanding. ChatMed-VHI addresses this need. In this work, we introduced ChatMed-VHI, an instruction-tuned NER model based on ChatGPT-3.5 turbo, designed to extract virus-host interactions (VHIs) from full-length biomedical research articles. ChatMed-VHI operates on extended context windows of 2-6 sentences, enabling it to model short-to mid-range cross-sentence dependencies, such as resolving co-references or capturing interactions that span multiple sentences. By curating training data from both narrative (*Results*, *Materials and Methods*) and non-narrative (tables in both main and supplementary materials) sections, we enable the model to capture contextual and structured interaction information more comprehensively. This improves extraction accuracy and supports the construction of more holistic and context-aware biomedical datasets.

Next, we systematically compared two classes of models for VHI extraction: supervised fine-tuned BERT-based models (PubMedBERT, BioLinkBERT) and instruction-tuned GPT-based models (ChatGPT-3.5). Our results demonstrated that instruction-tuned GPT-based models consistently outperform their BERT-based counterparts in low-resource settings. Specifically, ChatMed-VHI achieved an F1 score close to 90% with fewer than 500 training examples, significantly surpassing PubMedBERT under identical training conditions. These findings highlight the data-efficient, high-performance potential of instruction-tuned LLMs for biomedical named entity recognition.

In summary, we present a practical and scalable approach for extracting VHIs from diverse textual sources within biomedical literature. Our findings underscore the promise of instruction-tuned LLMs in enhancing biomedical information extraction, particularly in settings where training data is limited or distributed across complex document structures.

## Materials and Methods

The overall methodology for comparing fine-tuning approaches to extract virus-host interaction entities is illustrated in **Figure 1**. The workflow begins with collecting interaction data from full-text articles, including the *Results*, *Materials and Methods* sections, and both main-text and supplementary tables. The raw data are processed into training, development, and test sets. Two fine-tuning strategies were applied: (1) a span-based supervised fine-tuning approach using BERT-based models, and (2) an instruction-tuning approach using ChatGPT. Model performance was then evaluated on a held-out test set to assess their effectiveness in extracting virus-host interactions.

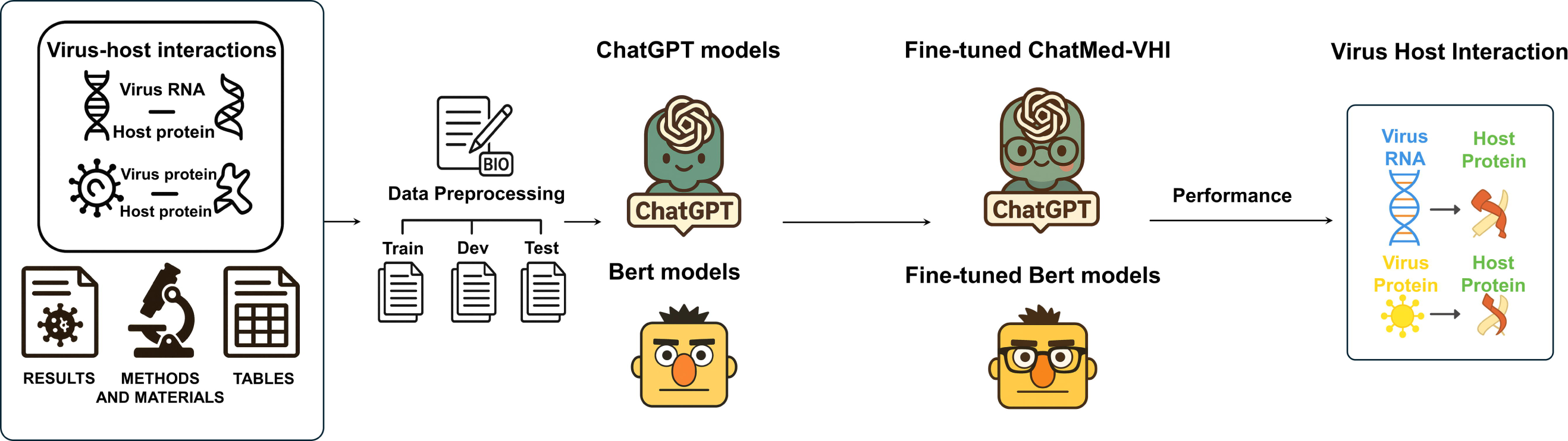

### Dataset Construction

This study focuses on NER for virus-host interactions. The curated dataset comprises three categories of information: (1) virus-specific entities, (2) host-related entities, and (3) experimental context (**Figure 2**). A full schema and description of each field are provided in **Table 1**. Data is extracted from three sources within PubMed articles: (1) *Results* sections (2) *Materials and Methods* sections, and (3) tables in both main and supplementary materials (**Figure 2**).

**Figure 2.**
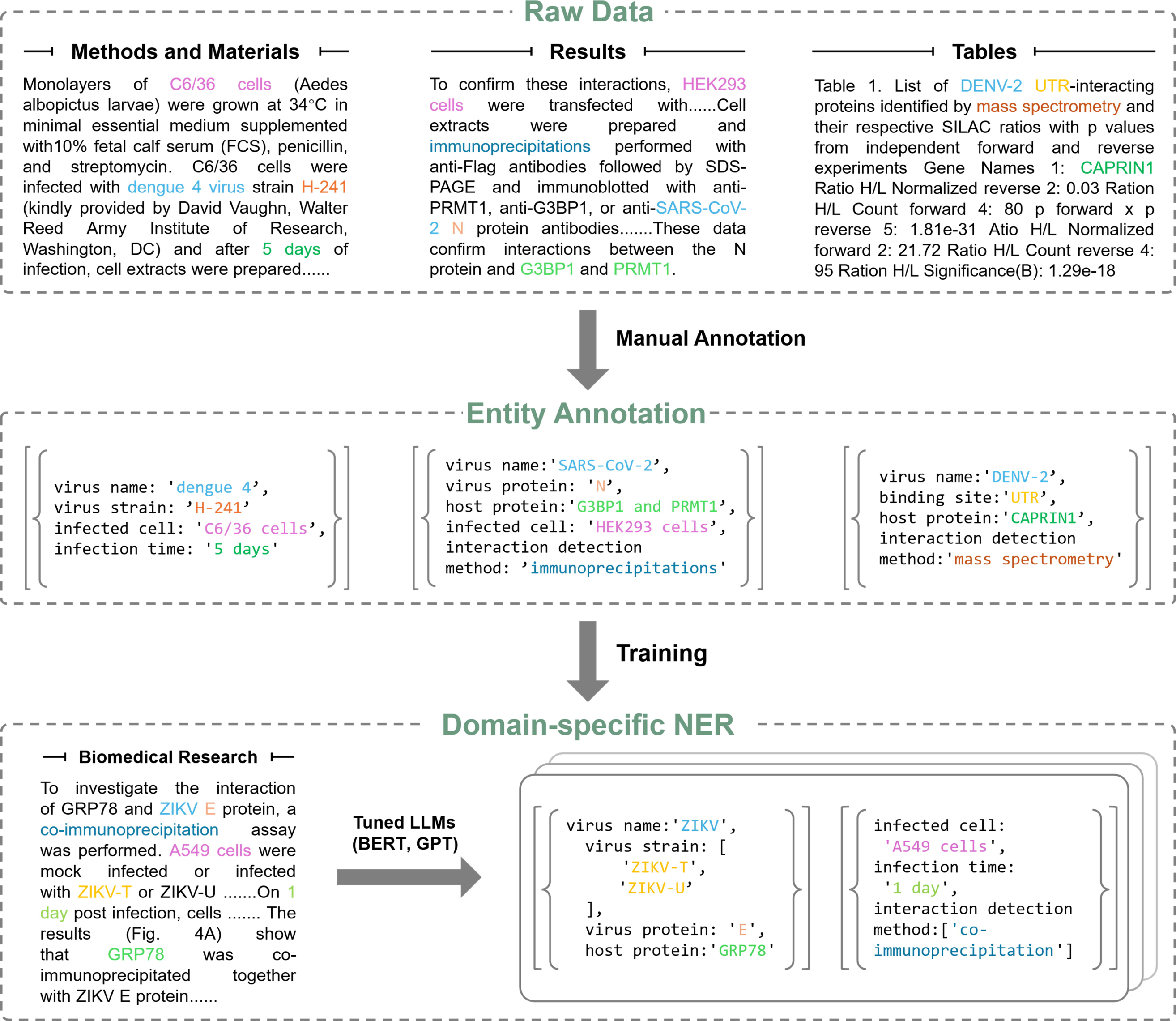
Curation of a comprehensive virus-host interaction dataset from full-text articles for large language model training.

| Field Name | Description |
| --- | --- |
| Virus Information |  |
| virus_name | Name of the virus (e.g., SARS-CoV-2) |
| virus_strain | Specific viral strain (e.g. B.1.1.7) |
| virus_protein | Viral protein involved in interaction (e.g., capsid) |
| RNA_structure_preference | RNA secondary structure preference (e.g., stem-loop) |
| binding_site | Viral binding site interacting with the host (e.g., UTR) |
| virus_function_on_host | Functional effect of the virus on the host system |
| Host Information |  |
| host_protein/RBP/host_factor | Host protein or RNA-binding protein involved (e.g., ACE2) |
| host_species | Host species (e.g., Homo sapiens, Mus musculus) |
| host_function_on_virus | Functional effect of the host factor on the virus |
| Interaction Context |  |
| interaction_detection_method | Experimental method used (e.g., CLIP, RIP, Y2H) |
| cell_type | Cell type used in infection experiments (e.g., HeLa cell) |
| cell_origin | Source/type of the cell (e.g., lung, kidney) |
| infection_time | Timepoint of infection (e.g., 12h, 24h post-infection) |

From the narrative text, we extracted sentences containing virus- and host-related entities and their interactions. In parallel, RPIs and PPIs were also collected from tables. For each record, we constructed natural language sentences by integrating table titles, column headers, and corresponding cell values. To prevent overfitting the model to a fixed format, the field-value pairs in each constructed sentence were randomly shuffled (**Figure 2**). These sentences were then processed in the same manner as narrative text.

### Data formatting for model fine-tuning

BioLinkBERT-base, PubMedBERT-base and ChatGPT-3.5 were fine-tuned on our curated virus-host interaction dataset.

For BERT-based models, data were formatted in a SQuAD-style question answering format. Each instance was stored as a JSON object containing three fields: *context* (the original sentences), *question* (the description of a target entity), and *answers* (a dictionary with the extracted text entity and its starting character index). An example is shown in **Figure 3A**.

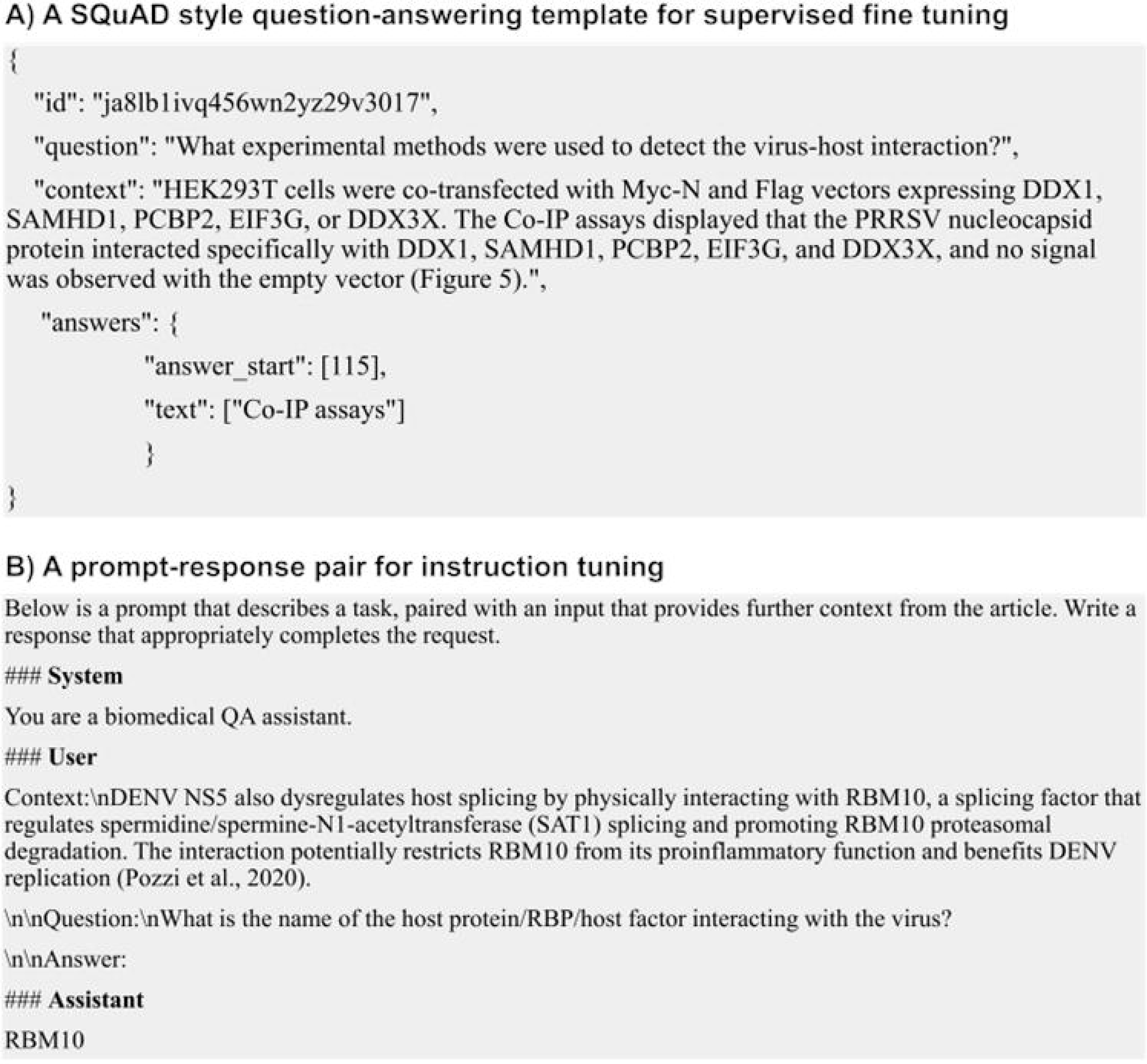

For ChatGPT instruction tuning, data were reformatted into a conversational prompt structure with three fields: *System* (the role or behavior of the assistant), *User* (a task description with the input context), *Assistant* (the target entity). Additional prompts were created by varying the instructions to extract different entity types (e.g., virus names, host proteins). An example is shown in **Figure 3B**.

### Instruction tuning of ChatGPT

Instruction tuning refers to training LLMs using datasets annotated via natural language instructions, improving zero-shot performance on unseen tasks [31]. This approach has been successfully applied to models such as LLaMA-1 7B, LLaMA-2 7B [30], Stanford Alpaca [32] and Flan-UL2 [33]. In this study, we applied instruction tuning to ChatGPT-3.5 via structured prompting. Since direct parameter updates are not possible through OpenAI’s API, fine-tuning is performed by training the model to generate target entities in response to structured instruction-context pairs. An instruction-response template is illustrated in **Figure 3B**.

### Supervised fine-tuning of BERT-based models

Supervised fine-tuning adapts a pretrained model to a specific downstream task by updating its parameters using labeled data and a task-specific supervised loss function. As described in the original BERT paper, a single output layer can be added to support fine-tuning across a range of tasks [15]. Supervised fine-tuning has been widely adopted and validated in biomedical models such as BioBERT [21], RoBERTa [34] and PubMedBERT [20]. In our study, we fine-tuned PubMedBERT-base and BioLinkBERT-base using the curated dataset formatted in the SQuAD-style question-answering template. The model was trained to predict start and end tokens corresponding to the correct entity span in each sentence. An example of the SQuAD style question-answering template is illustrated in **Figure 3A**.

### Baselines

#### PubMedBERT

PubMedBERT [20] is a domain-specific language model pretrained on abstracts and full-text articles from PubMed and PubMed Central. In this study, we fine-tuned PubMedBERT on our curated virus-host interaction dataset using standard training and validation splits. Hyperparameter tuning was performed using the following search space: learning rate∈{1×10^−5^, 3×10^−5^, 5×10^−5^}, sequence length∈{256, 384, 512}, batch size∈{16, 32}, number of epochs ∈{15, 20, 25}, and a fixed dropout rate of 0.1.

#### BioLinkBERT

BioLinkBERT [22] extends PubMedBERT by incorporating citation links between articles during pretraining, allowing it to capture both textual and relational information. This model surpassed the previous pretrained model PubMedBERT [20], and achieved better performance on the BLURB benchmark dataset [22]. For consistency, the same hyperparameter grid was applied during BioLinkBERT fine-tuning.

#### GPT Fine-tuning

We fine-tuned gpt-3.5-turbo-0125 using OpenAI’s API with instruction-style prompt-response pairs. The final model, ft:gpt-3.5-turbo-0125:personal::Balpe7G4, was trained on 1.76M tokens from train_finetune_cleaned.jsonl, validated on dev_finetune_format.jsonl. Fine-tuning was supervised, with epochs (3), batch size (8), learning rate multiplier (2), and seed (1161655790). The job (ftjob-6U5kqWwzikPK0IrdQzhuwQjK) completed on May 24, 2025. Validation loss was 0.178. GPT-3.5 was chosen for its tunability and lower inference cost. GPT-4, while more capable, did not support fine-tuning via the API at the time of this work. An example prompt we used for instruction tuning is illustrated in **Figure 3B**.

#### GPT Zero-shot/Few-shot

To further evaluate model generalization, we assessed GPT-3.5 (gpt-3.5-turbo-0125) and GPT-4o (gpt-4o-2024-08-06) on the test sets using both zero-shot and few-shot settings. For five-shot learning, five randomly selected training examples were included in the prompt. The prompt format remained consistent with that used in instruction tuning (**Figure 3B**).

### Evaluation Metrics

Model performance was assessed using precision, recall, and F1 score. We employed a strict (exact match) criterion, where predicted entities were considered correct only if both their span and label exactly matched the gold-standard annotations. For GPT-based models (both zero-shot/few-shot and fine-tuned), this exact match evaluation may underestimate true performance, as even minor deviations from the reference span (e.g., punctuation, partial boundary mismatches) are considered incorrect. To address this, we also report partial match results in the *Results* section, based on the overlap between predicted and gold spans.

## Results

The virus-host interaction dataset constructed for model fine-tuning consists of 5,328 instances derived from 225 biomedical articles. Specifically, the dataset includes 3,395 instances of virus-host PPIs and 674 instances of virus-host RPIs, extracted from the *Results* and *Materials and Methods* sections. An additional 1,259 instances, comprising 566 PPIs and 793 RPIs, were obtained from tables in both the main text and supplementary materials. Each table entry was converted into a natural language sentence and annotated following the same schema (see *Materials and Methods*). The test set consists of 534 instances, including 234 mentions of virus-related entities, 163 mentions of host-related entities, and 137 mentions of experimental context (see **Figure 2**).

### Model Performance

The performance of BERT-based and ChatGPT models on the test set is summarized in **Table 2**. Using a consistent prompt structure (**Figure 3B**) across zero-shot, five-shot, and fine-tuned settings, we found that ChatMed-VHI performed comparably to, and in some cases outperformed, traditional BERT-based models. Specifically, ChatMed-VHI shows a slightly higher F1 score (0.9223) than fine-tuned PubMedBERT (0.9104), although the difference was not statistically significant. Despite incorporating citation links in pretraining, fine-tuned BioLinkBERT achieved slightly lower precision (0.9133 vs. 0.9229) and F1 score (0.9062 vs. 0.9104) than fine-tuned PubMedBERT. Both fine-tuned BERT-based and GPT-based models outperformed zero-shot and five-shot GPT models. Among few-shot settings, ChatGPT-3.5 and ChatGPT-4o showed comparable performance, with F1 scores of 0.8074 and 0.8027, respectively. In the zero-shot setting, ChatGPT-4o outperformed ChatGPT-3.5 (F1: 0.7913 vs. 0.7088), demonstrating its superior generalization ability in the absence of task-specific training.

| Model | Metrics | Strict match | Partial match |
| --- | --- | --- | --- |
| PubMedBERT(fine-tuned) | P | **0.9229** | 0.9536 |
|  | R | 0.8917 | 0.9490 |
|  | F1 | **0.9104** | 0.9425 |
| BioLinkBERT(fine-tuned) | P | **0.9133** | 0.9523 |
|  | R | 0.8991 | 0.9566 |
|  | F1 | **0.9062** | 0.9445 |
| ChatMed-VHI | P | **0.9133** | 0.9549 |
|  | R | 0.9336 | 0.9662 |
|  | F1 | **0.9223** | 0.9544 |
| Chatgpt-4o(zero-shot) | P | 0.8418 | 0.8476 |
|  | R | 0.7465 | 0.8615 |
|  | F1 | 0.7913 | 0.8266 |
| Chatgpt-4o(five-shot) | P | 0.8257 | 0.8376 |
|  | R | 0.7810 | 0.8625 |
|  | F1 | **0.8027** | 0.8300 |
| Chatgpt-3.5(zero-shot) | P | 0.7450 | 0.7638 |
|  | R | 0.6760 | 0.7766 |
|  | F1 | 0.7088 | 0.7423 |
| Chatgpt-3.5(five-shot) | P | 0.8320 | 0.8385 |
|  | R | 0.7842 | 0.8650 |
|  | F1 | **0.8074** | 0.8309 |

### Prediction Error Analysis

Model performance was further analyzed using both strict match and partial match criteria. A strict match requires the predicted span to align perfectly with the gold-standard annotation, whereas partial match allows for token-level overlap between predicted and gold spans. As shown in **Table 2**, partial match scoring consistently resulted in higher F1 values across all models. For example, the F1 score of fine-tuned PubMedBERT increased from 0.9104 (strict) to 0.9425 (partial), and ChatGPT-3.5 improved from 0.9223 (strict) to 0.9544 (partial). The largest improvements under partial matching were observed in ChatGPT-3.5 model (zero-shot, 4.73% increase), followed by ChatGPT-4o model (zero-shot, 4.46% increase).

Manual error analysis reveals three main error types, following an established taxonomy [35]: (1) missing entities, where the model failed to predict any relevant entity; (2) wrong entities, where incorrect entities were predicted by the model; (3) boundary issues, where the entity type was correct but the span did not match the gold standard. As shown in **Figure 4A**, boundary issues accounted for the majority of errors across all models. Fine-tuned GPT-3.5 produced the fewest total errors (45 instances), including 15 wrong entities and 30 boundary issues. BioLinkBERT and PubMedBERT yielded 33 and 37 boundary issues, respectively, while GPT-3.5 and GPT-4o (five-shot) exhibited the highest total error counts.

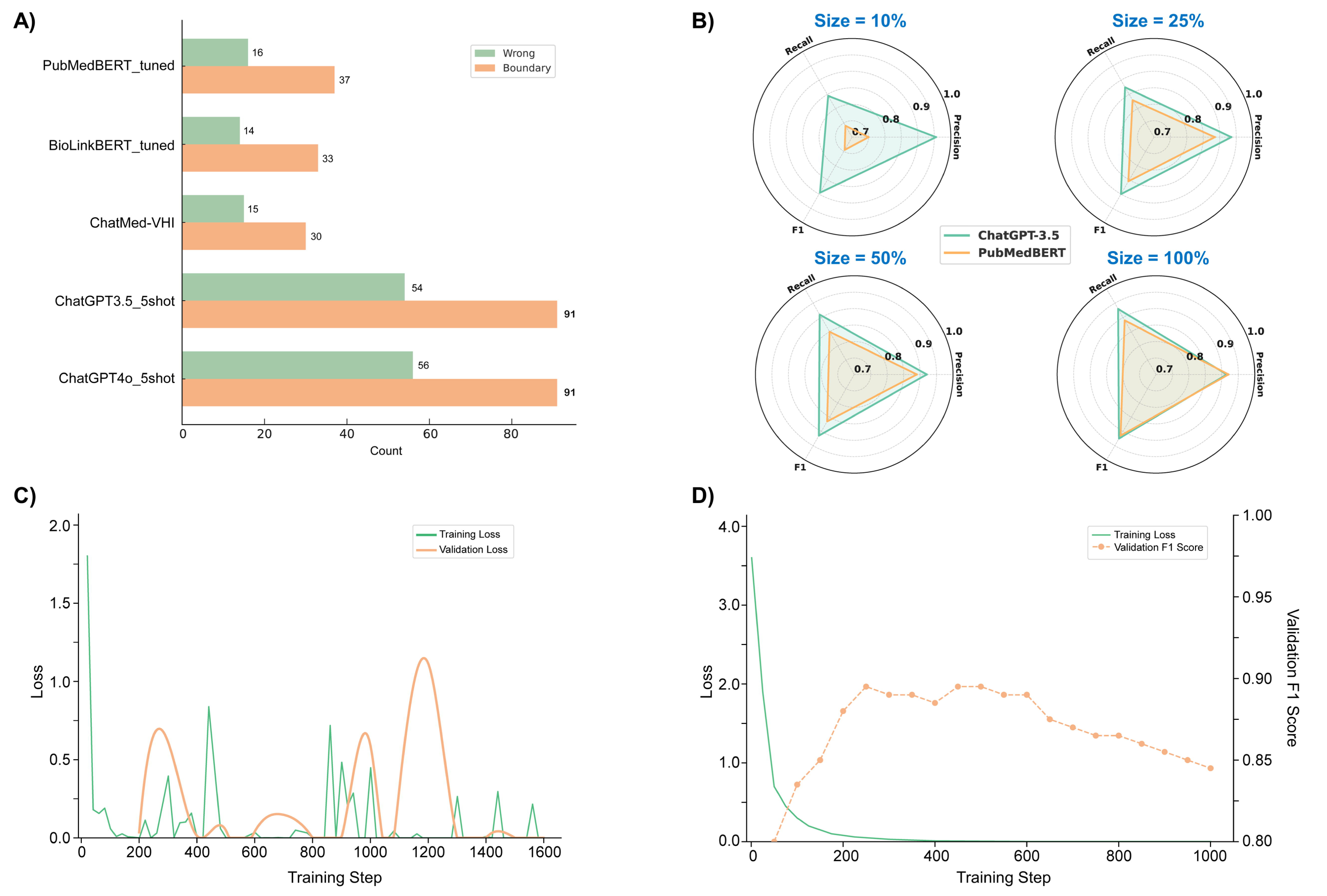

Further analysis highlighted model limitations in handling multi-entity spans. For example, when prompted to extract all host proteins involved in viral interactions, the correct answer “ANPEP, RAD18, APEX, POLH, APEX1, TERF2, RAD51, CDC7, USP7, XRCC5, FEN1, PCNA” was treated as a coordinated entity group, labeled with a BIO sequence of ‘**B I O B I O B I O B I O B’**. However, PubMedBERT only correctly recognized “ANPEP” (tagged as ‘**B O O O O O O O O O O O’**), missing the rest, whereas GPT-3.5 successfully recovered the entire set with near-perfect BIO alignment.

Another common error involved abbreviation mismatches. In response to the query “Which viral protein interacts with the host?”, the gold label was “capsid” (B). However, both PubMedBERT and GPT-3.5 incorrectly predicted “CA” (B), a commonly used abbreviation for HIV capsid protein. Although the predicted span followed the correct BIO structure, the semantic mismatch caused evaluation failures under the strict matching criterion. Similar abbreviation-related errors were also observed in GPT-3.5 and GPT-4o under both zero-shot and few-shot settings.

### Effect of dataset size on LLM performance

We next examined the effect of training dataset size on model performance, as shown in **Figure 4B** and **Table 3**. To simulate low-resource scenarios, the dataset was reduced to 50% (2,131 instances), 25% (1,065 instances), and 10% (426 instances) of its original size. The performance of ChatGPT-3.5 decreased only modestly across these reductions. When using 50% data, the F1 score dropped by just 0.77%. With 25% and 10% data, the F1 scores declined by 2.33% and 2.79%, respectively. Precision also decreased, from 0.953 with the full dataset to 0.913 when trained with only 10% data. These results indicate that ChatGPT-3.5 maintains relatively strong performance even under limited supervision.

| Model |  | Training Size |  |  |  |
| --- | --- | --- | --- | --- | --- |
|  |  | 10% | 25% | 50% | 100% |
| PubMedBERT | Strict | 0.751/0.742/**0.746** | 0.885/0.829/**0.856** | 0.889/0.852/**0.870** | 0.923/0.892/**0.910** |
|  | Partial | 0.849/0.851/0.823 | 0.926/0.917/0.910 | 0.937/0.933/0.933 | 0.954/0.949/0.943 |
| ChatGPT3.5 | Strict | 0.953/0.847/**0.897** | 0.935/0.869/**0.901** | 0.921/0.909/**0.915** | 0.913/0.934/**0.922** |
|  | Partial | 0.949/0.929/0.927 | 0.949/0.939/0.936 | 0.955/0.957/0.949 | 0.955/0.966/0.955 |
| Numbers in bold represents the best performance for each dataset. |  |  |  |  |  |

In contrast, PubMedBERT showed a more substantial decline in performance as the dataset size decreased. Compared to ChatGPT-3.5, the F1 score of PubMedBERT dropped by 4.60% with 50% data, 6.31% with 25% data, and 21.98% with 10% data. ChatGPT-3.5 consistently outperformed PubMedBERT across all training sizes. Moreover, the performance gap between the two models widened with decreasing data availability. With the full dataset, ChatGPT-3.5 outperformed PubMedBERT by 1.32%. This margin increased to 5.17% with 50 % data, 5.26% with 25% data, and reached its maximum at 20.25%, when only 10% training data was used.

These performance trends were further reflected in the training dynamics of ChatGPT-3.5 and PubMedBERT, illustrated in **Figure 4C** and **4D**. ChatGPT-3.5 showed a rapid reduction in training loss during the initial phase, decreasing from approximately 1.8 to below 0.2 within the first 100 steps. However, its validation loss exhibited considerable fluctuations throughout training, with sharp peaks occurring around steps 250, 1000, and 1200. These oscillations suggest unstable generalization behavior under the instruction-tuning framework. In contrast, PubMedBERT displayed a smooth and consistent decline in training loss, from 3.6 to nearly zero by step 1000. The validation F1 score for PubMedBERT increased dramatically in the first 200 steps, reaching approximately 0.91. However, as training continued, this score gradually declined to 0.85, indicating that PubMedBERT generalized well initially but experienced mild overfitting in later epochs.

In contrast to prior instruction-tuned or prompt-based frameworks such as BioInstruct [29] and ChatExtract [12], our ChatMed-VHI system provides several key advantages in the context of virus-host interaction extraction. BioInstruct, built upon LLaMA and trained on synthetic biomedical instructions, demonstrated strong general performance across various biomedical NLP tasks [29]. However, it was not optimized for full-text processing or section-aware extraction, and its evaluations primarily focused on short inputs and standard benchmarks rather than complex entity structures such as VHIs. Similarly, ChatExtract leverages conversational prompting to achieve high precision and recall in zero- or few-shot settings [12], but it was originally designed for structured numerical data in materials science and relies on sentence-level context without instruction fine-tuning. This makes it less effective in capturing cross-sentence dependencies, resolving abbreviations and synonyms, or integrating section-specific cues,all of which are crucial in VHI tasks.

In contrast, ChatMed-VHI is instruction-tuned on full-text biomedical articles that include both narrative (e.g., Results, Methods) and non-narrative (e.g., tables, supplementary) content. It is explicitly optimized to extract complex and variable-length biomedical entities under weak supervision, achieving near-human performance (F1: ∼90%) with only 426 training examples. By combining domain-adapted instructions, section-aware input windows, and carefully engineered prompt constraints, ChatMed-VHI demonstrates stronger generalizability, contextual robustness, and data efficiency compared to previous tools.

## Discussion

In this study, we proposed a data-efficient approach to extract PPIs and RPIs from all data-rich sections of biomedical articles and systematically compared fine-tuning strategies using ChatGPT and BERT-based models. A key finding of our study is the markedly different training dynamics exhibited by ChatGPT-3.5 and PubMedBERT under varying training data sizes (**Table 3**). For BERT-based models, all performance metrics (F1, precision, and recall) improved steadily with increasing training data, ultimately reaching a strong precision score of 0.923. In contrast, ChatGPT-3.5 exhibited diminishing returns, where adding more training data beyond a certain point did not lead to further improvements [36]. Notably, its precision decreased from 0.953 with 10% training data to 0.913 with the full dataset. A similar pattern was reported in a study on LLaMA, where fine-tuning led to a decline in performance relative to the baseline model [37].

The training dynamics illustrated in **Figure 4C** and **4D** offer insight into the divergent performance trends. ChatGPT-3.5 exhibits unstable learning dynamics, with rapid reduction in training loss during the first 100 steps (from ∼1.8 to <0.2), followed by significant oscillations in validation loss throughout training. This instability may be attributed to overfitting to the fixed prompt structure, causing the model to generate templated responses that diverge from the gold-standard answer spans. Such deviations can lead to higher rates of span mismatches and false positives during evaluation.

Interestingly, despite the decline in precision, ChatGPT-3.5’s overall F1 score improved with larger training datasets due to substantial gains in recall. This indicates the model became more effective at identifying correct answers and reducing the number of false negatives, even though the outputs were less tightly aligned with the gold annotations. According to OpenAI’s official guidance (https://platform.openai.com/docs/guides/fine-tuning), there is no recommended number of training examples as the optimal dataset size is often case-dependent. However, it is generally advised to begin with 50 well-crafted examples, as performance improvement typically occurs within the range of 50-100 examples. In our case, ChatGPT-3.5 achieved an F1 score near 90% using only 426 examples, underscoring its efficiency and strong performance in domain-specific tasks with limited supervision. In contrast, PubMedBERT demonstrated more stable and interpretable learning. Its training loss declined steadily from 3.6 to near zero, while its validation F1 score peaked at around step 200 before gradually declining, consistent with mild overfitting. This robust and consistent improvement is likely due to BERT’s architecture being well-suited for token-level classification tasks, reinforcing the reliability of traditional BERT models when scaled with more annotated data.

As illustrated in **Figure 4A**, BERT-based models underperformed ChatGPT-3.5 in recognizing coordinated or list-like mentions (e.g., multiple host proteins in a single sentence). This is largely due to BERT’s reliance on identifying contiguous spans, making it less effective at extracting entities dispersed across non-adjacent text segments. Although newer variants such as SpanBERT[38] address this issue to some extent, they still struggle with complex, nested biomedical sentences. We also found that well-designed prompts could improve ChatGPT’s performance. Without restrictive instructions, ChatGPT tends to generate unnecessary long responses or reformulated the input. To mitigate this, we added explicit constraints in the prompt, such as “Extract the answer to the following question using only the words from the given context” (**Figure 3B**), which led to more concise and context-aligned outputs. To further support accurate extraction, we also incorporated contextual cues and synonymic variations into the prompt design (**Table 1**). For instance, RNA-binding proteins (RBPs) may be referred to as “cell protein”, “host factor” or “cellular factor”, while tissue-related entities may appear under terms like “histology”, “tissue”, or “organ”.

Despite these improvements, extraction accuracy was still affected by inconsistent biomedical nomenclature (**Figure 4A**). Prediction errors often stemmed from lexical variability, such as “human angiotensin-converting enzyme” versus “hACE”, or inconsistent expression of time (e.g., “24 hpi” vs. “24 hours”). These semantically equivalent but lexically distinct terms reduced precision under strict evaluation. Addressing this challenge through entity normalization or integration with biomedical ontologies remains a key direction for improving NER performance in biomedical applications

Finally, as reported in **Table 2**, we observed that adding five-shot demonstrations unexpectedly reduced ChatGPT-4o’s precision. We attribute this to two primary factors: (1) the limited number of examples (n = 5) was insufficient to cover the broad diversity of query types in our task; and (2) the models exhibited signs of prompt-style overfitting. In contrast, zero-shot prompting leveraged ChatGPT-4o’s inherent generalization capacity, yielding more robust performance across varied question types. The narrowing performance gap between ChatGPT-4o and 3.5-turbo under the five-shot setting further suggests that weaker models benefit more from few-shot guidance, while stronger models may already operate near their task-specific performance ceiling.

Compared to prior instruction-tuned models such as BioInstruct [29], which were trained on generalized biomedical NER tasks, ChatMed-VHI was explicitly designed for extracting virus-host interaction entities from full-length biomedical articles. While BioInstruct demonstrated strong performance on benchmark datasets, its training did not include full-text documents or VHI-specific contexts, limiting its applicability to complex multi-sentence interaction mentions. Likewise, prompt-based systems like ChatExtract [12] have shown that conversational LLMs can extract structured data from full text without fine-tuning. However, ChatExtract focuses primarily on materials science tasks and tabular data, relying on zero- or few-shot prompts that lack adaptability to biomedical domains with diverse and complex synonyms, and coreference patterns like virus-host biomolecular interaction.

In contrast, ChatMed-VHI integrates instruction tuning with full-text supervision, enabling it to handle dispersed, synonym-rich biomedical entities that span multiple sentences and sections. Our model further benefits from structured prompt design, synonym-aware labeling, and section-specific training examples, resulting in robust performance even under low-resource settings. This makes ChatMed-VHI a domain-specialized, scalable solution for high-accuracy biomedical entity recognition beyond abstract-level input.

## Conclusion

We systematically evaluated fine-tuning frameworks for extracting virus-host biomolecular interactions. As part of this work, we introduced ChatMed-VHI, an instruction-tuned variant of ChatGPT-3.5, tailored for recognizing virus–host interaction entities. To support a comprehensive evaluation, we constructed a rich, domain-specific dataset from full-text biomedical articles, including content from the *Materials and Methods* sections and tables in both main and supplementary materials. we used this dataset to compare two fine-tuning approaches: supervised fine-tuning of BERT-based models (PubMedBERT, BioLinkBERT) and instruction tuning of ChatGPT-3.5. Our results revealed that ChatMed-VHI significantly outperformed PubMedBERT under low-resource conditions (less than 500 training examples). The significance of this work is twofold. Firstly, our full-text curation methodology provides a novel approach for creating more context-rich and holistic biomedical datasets that better reflect the complexity of real-world scientific literature. Secondly, ChatMed-VHI demonstrates the effectiveness and robustness of instruction tuning for creating a high-performance discipline-specific NLP tool, especially when annotated data are scarce. This finding opens new avenues for future work. An immediate action could be extending this framework from entity recognition to relation extraction, ultimately enabling the construction of a comprehensive virus-host biomolecular interactions knowledge graph.

## Code Availability

The source code and datasets used in this study are available at https://github.com/benkyusimasu/ChatMed-VHI.

## Acknowledgments

We thank our colleagues and reviewers for their constructive comments and insights that greatly improved the manuscript. The authors acknowledge financial support from the National Natural Science Foundation of China (NSFC) under Grant No. 23CAA01280 (Outstanding Young Scientists Fund – Overseas) and Grant No. 22307122 (Youth Fund Project), as well as from the Chinese Academy of Sciences (CAS), Hangzhou Institute of Medicine, and Zhejiang Cancer Hospital.

## Author Contributions

Z.Z. conceived the core idea of the study, led the data collection and curation process, and contributed to the design of the NLP system and model code. He also drafted the initial version of the manuscript.

W. L designed the code and model architecture, and co-authored the manuscript. He provided key insights into BioNLP from his expertise in question answering and information extraction.

Y.C. assisted with the data collection and curation process.

J.R., B.H., Q.S. participated in the formulation, discussion, and writing of the project, offering valuable suggestions from a biomedical research perspective.

H.W. is the planner of the whole project. She was responsible for writing the proposal for the project, planning the direction and progress of the whole project, discussing and writing the final paper.

